# Physiological assembly of functionally active 30S ribosomal subunits from *in vitro* synthesized parts

**DOI:** 10.1101/137745

**Authors:** Jun Li, Brook Wassie, George M. Church

**Affiliations:** Department of Genetics, Harvard Medical School, 77 Avenue Louis Pasteur, Boston, MA 02115, USA; Department of Biological Engineering, Massachusetts Institute of Technology, Cambridge, MA 02139, USA

## Abstract

Synthetic ribosomes *in vitro* can facilitate engineering translation of novel polymers, identifying ribosome biogenesis central components, and paving the road to constructing replicating systems from defined biochemical components. Here, we report functional synthetic *Escherichia coli* 30S ribosomal subunits constructed using a defined, purified cell free system under physiological conditions. We test hypotheses about key components of natural ribosome biogenesis pathway as required for efficient function – including integration of 16S rRNA modification, cofactors facilitated ribosome assembly and protein synthesis in the same compartment *in vitro*. We observe ~17% efficiency for fully synthetic 30S and ~70% efficiency from *in vitro* transcribed 16S rRNA assembled with natural proteins. We observe up to 5 fold improvement over previous crude extracts. We suggest extending the minimal list of components required for central-dogma replication from the 151 gene products previously reported to at least 180 to allow the speed and accuracy of macromolecular synthesis to approach native *E. coli* values.

## INTRODUCTION

The *Escherichia coli* 70S ribosome is a complex ribonucleoprotein machine composed of two subunits: 30S and 50S. The 30S subunit contains a 16S rRNA and 21 ribosomal proteins (r-proteins), whereas the 50S subunit contains two RNAs, 5S rRNA and 23S rRNA, and 33 r-proteins (Supplementary Table S1).

*E. coli* ribosome reconstitution or assembly from purified native components into functionally active 30S and 50S subunits was first achieved under non-physiological conditions ~40 years ago. The conventional 30S subunit reconstitution protocol involves a one-step incubation at 20 mM Mg^2+^ and 40°C (1), while the one for 50S subunit involves a non-physiological two-step high-temperature incubation, first at 4 mM Mg^2+^ and 44°C, then at 20 mM Mg^2+^ and 50°C (2) (Figure 1a). These non-physiological conditions preclude coupling of ribosome assembly and protein synthesis in one compartment. Meanwhile, conventionally reconstituted 30S and 50S subunits from natural proteins and *in vitro* transcribed rRNAs lacking post-transcriptional modifications are ~50% (3) and 4 orders of magnitude (4) less active respectively compared to native ribosomes.

**Figure 1.**
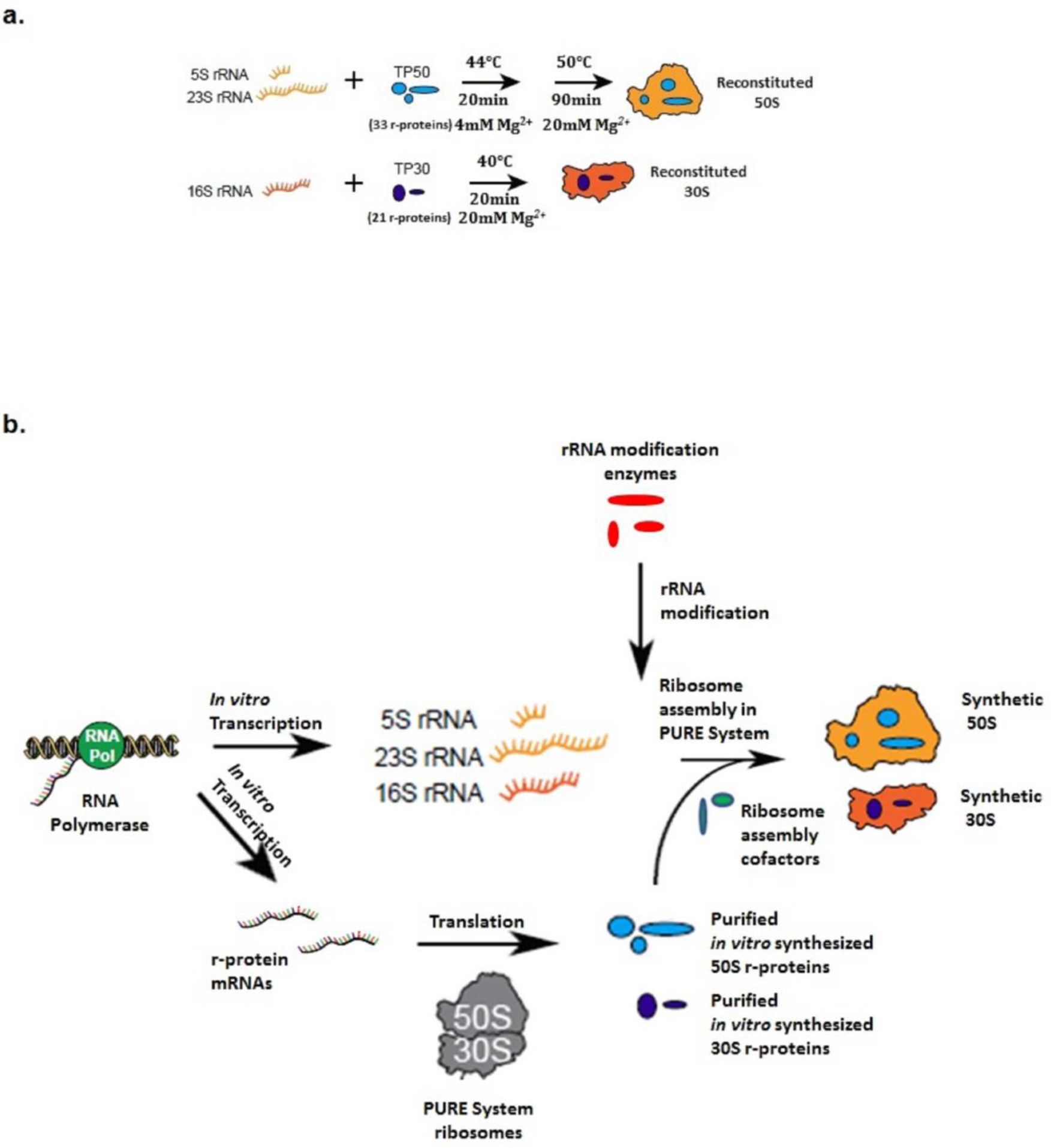
Conventional and our integrated method to construct *E. coli* ribosomes *in vitro*. **(a)**. Conventional method for ribosome reconstitution. **(b).** Our integrated method to construct synthetic ribosomes in PURE Δ ribosome system under physiological conditions by supplementing assembly cofactors and rRNA modification enzymes.

By contrast, *E. coli* ribosome biogenesis *in vivo* is a highly coordinated process involving many protein factors, such as endonucleases, rRNA and r-protein modification enzymes, GTPases and helicases (5, 6). RimM, RimN, RimP, RbfA, Era and RsgA are believed to associate with 30S subunit as assembly cofactors (Table 1), while CsdA, DbpA, Der, and SrmB are supposed to associate with 50S subunit assembly (Supplementary Table S2). To date, all of the eleven 16S rRNA modification enzymes have been identified (Table 2), while two of the 23S rRNA modification enzymes on position 2449 and 2501 essential for 50S subunit function remain to be unknown (7) (Supplementary Table S3).

**Table 1.** Summary of 30S ribosome assembly cofactors

| **Factor** | **Possible functions** | **Reference** |
| --- | --- | --- |
| Era | Era is a GTPase that binds to (pre-) 30S subunit to facilitate processing of the 3' end of the 16S rRNA precursor. | (14) |
| RimM | RimM binds to the head of the small ribosomal subunit to facilitate assembly. | (29,30) |
| RimN | RimN has been suggested to bind the 16S rRNA to promote proper processing. | (31) |
| RimP | RimP binds to the 16S rRNA, helps r-proteins bind faster and is thought to act towards the end of 30S assembly. | (32) |
| RfbA | RfbA binds to 30S subunit to facilitate subunit assembly. Overexpression suppresses a cold sensitive C23U mutation in the 16S rRNA. | (33) |
| RsgA | RsgA has a putative role in small subunit ribosomal assembly. | (34) |

**Table 2.** Modified nucleotides of *E. coli* 16S rRNA and their modification enzymes

| **Nucleotide** | **Modification** | **Gene** | **Substrate** | **Reference** |
| --- | --- | --- | --- | --- |
| 516 | $\Psi$ | RusA (yejD) | Semi-assembled 30S | (35) |
| 527 | $m^7G$ | RsmG (gidB) | Semi-assembled 30S | (36) |
| 966 | $m^2G$ | RsmD (yhhF) | 30S | (37) |
| 967 | $m^5C$ | RsmB (fmu) | 16S rRNA | (38) |
| 1207 | $m^2G$ | RsmC (yjjT) | Semi-assembled 30S | (39) |
| 1402 | $m^4Cm$ | RsmH (yabC),<br>RsmI (yraL) | 30S | (40,41) |
| 1407 | $m^5C$ | RsmF (yebU) | 30S | (42) |
| 1498 | $m^3U$ | RsmE (yggJ) | 30S | (43) |
| 1516 | $m^2G$ | RsmJ (yhiQ) | 30S | (44) |
| 1518 | $m^6_2A$ | RsmA (ksgA) | Semi-assembled 30S | (45) |
| 1519 | $m^6_2A$ | RsmA (ksgA) | Semi-assembled 30S | (45) |

By mimicking chemical conditions in the cytoplasm, synthetic ribosomes can be assembled from *in vitro* transcribed rRNAs and natural proteins in crude *E. coli* S150 extracts under physiological conditions (8). However this system is undefined and it is hard to know which factors in the extracts facilitate ribosome assembly or modify the rRNAs. In this study, we use a well-defined reconstituted cell-free protein synthesis system, the PURE system lacking ribosomes (PURE Δ ribosome system) (9), to construct synthetic *E. coli* ribosomes under physiological conditions. Our approach attempts to mimic ribosome biogenesis pathway *in vivo* by supplementing ribosome assembly cofactors, rRNA modification enzymes, allowing rRNA modification, cofactor facilitated ribosome assembly, *in vitro* transcription and translation of a reporter gene to take place in the same compartment (Figure 1b). Since modification enzymes for two of the essential modifications on 23S rRNA are still unknown, we took 30S subunit as the subject to test our approach. Firefly luciferase (Fluc) was used as a reporter for the measurement of ribosome activity. We show that using our method, 30S subunits can be reconstituted under physiological conditions from natural components with 90% activity of native ribosomes by supplementing 30S ribosome assembly cofactors RimM, RimN, RimP, RbfA, Era and RsgA. We also show that by supplementing 30S ribosome assembly cofactors and the eleven 16S rRNA modification enzymes, semi-synthetic 30S can be reconstituted from *in vitro* transcribed 16S rRNA and natural r-proteins of 30S subunit (TP30) with ~70% activity relative to native ribosomes. Moreover, we demonstrate that using our method, fully synthetic 30S ribosomes can be constructed from *in vitro* synthesized 30S r-proteins and *in vitro* transcribed 16S rRNA with 17% activity compared to native ribosomes. We also demonstrate the synthesis of 50S r-proteins and the co-synthesis of 30S and 50S r-proteins in the PURE system. Therefore our integrated method can potentially be applied to construct fully synthetic 50S ribosomes and study the roles of 50S ribosome assembly cofactors and 23S rRNA modification enzymes in ribosome biogenesis. As a novel approach for synthetic ribosome construction and for studying ribosome biogenesis *in vitro* (10), our method enables the identification of essential ribosome biogenesis factors, solves critical barriers to construct synthetic ribosomes and synthetic life and also opens the door to build a protein based replicating system (11, 12).

## MATERIALS AND METHODS

### Molecular cloning

The sequences of yejD, gidB, fmu, yjjT, yabC, yraL, yebU, ksgA, era, rimM, rimN, rimP, rbfA and rsgA were amplified by PCR from *E. coli* MG1655 genomic DNA (ATCC) and the PCR products were inserted into NdeI and XhoI restriction sites of plasmid pET-24b (Novagen). Primers for cloning and affinity tag positions were listed in Supplementary Table S4 and S5. PET-15b yhhF is a gift from Dr. Andrzej Joachimiak, Argonne National Laboratory; pET-32a yggJ is a gift from Dr. Murray Deutscher, Department of Biochemistry and Molecular Biology, University of Miami Miller School of Medicine; pET-28a yhiQ is a gift from Dr. Kenneth E. Rudd, Department of Biochemistry and Molecular Biology, University of Miami Miller School of Medicine. The Fluc gene (from pZE21-luc (13)) was amplified by PCR and subcloned into pIVEX 2.3d (Roche) with restriction enzyme NcoI and XhoI. Sanger sequencing verification of each clone was performed by Genewiz.

Each 50S and 30S r-protein gene was PCR amplified from *E. coli* MG1655 genome (ATCC) with primers listed in Supplementary Table S6 and S7 and cloned to pET-24b (Novagen) using restriction enzyme NdeI and XhoI except for rplB. RplB was cloned to pIVEX 2.3d vector (Roche) with restriction enzyme NcoI and XhoI due to its internal cleavage site of NdeI. Each gene was cloned in its natural form with no additional amino acid or affinity tags. Sanger sequencing verification of clones was performed by Genewiz.

### Protein expression and purification

Eleven 16S rRNA modification enzyme constructs and pET-24b rimM, pET-24b rimN, pET-24b rimP, pET-24b rfbA, pET-24b rsgA were transformed into *E. coli* BL21/DE3 cells. Each transformed strain was cultured at 37°C in LB broth containing proper antibiotics to an A600 of 0.6. Isopropyl-b-D-thiogalactopyranoside at 1 mM was added to the culture, and incubation continued at 37 °C to an A600 of 1.4 –1.8. Cells were recovered by centrifugation and frozen at -70°C. Cell pellets were washed by binding buffer (20 mM sodium phosphate, 0.5 M NaCl, 40 mM imidazole, 6 mM β-mercaptoethanol pH 7.4) 3 times and lysed by BugBuster Protein Extraction Reagent (Novagen). Cell lysates were centrifuged at 15,000g for 25 min. Supernatant was applied to a GE histrap-Gravity column. His-tagged enzymes were eluted down by elution buffer (20 mM sodium phosphate, 0.5 M NaCl, 500 mM imidazole, 6 mM β-mercaptoethanol pH 7.4). Enzymes were then dialyzed to a storage buffer containing 50 mM Tris–HCl pH 7.6, 100 mM KCl, 1 mM DTT and 30% glycerol. PET-24b era was expressed in the PURE system due to the degradation problem when expressed *in vivo* (14). A 200 μl PURE system reaction was set up for Era. 200 μl Strep-Tactin magnetic beads (Qiagen) were washed three times with 1 ml wash buffer (50 mM Tris HCl pH 7.6, 0.5 M NaCl, 6 mM β-mercaptoethanol), resuspended in 200 μL wash buffer and then mixed with 200 μl PURE reaction on a rotator at 4°C for 3 hr. The beads were then immobilized with a magnet. Supernatant was discarded and beads were washed twice with 200 μl wash buffer. Strep-tagged Era was eluted by 150 μl elution buffer (50 mM Tris HCl pH 7.6, 0.5 M NaCl, 10 mM biotin, 6 mM β-mercaptoethanol), concentrated by EMD Amicon Ultracel 0.5 ml-3 K spin concentrator and exchanged to a storage buffer to a final concentration of 50 mM Tris-HCl pH 7.6, 100 mM KCl, 1 mM DTT and 30% glycerol. Purified enzymes were analyzed on 4-12% Bis-Tris Gel (Life Technologies) and concentrations were determined by Bradford Assay (Bio-Rad).

### Isolation of tightly coupled 70S ribosomes and ribosomal subunits

Tightly coupled 70S ribosomes, 30S and 50S subunits were prepared as described by Michael C Jewett (8). Results from three independent ribosome preparations and subsequent rRNA and total protein preparations were used and averaged to generate the final results shown in the manuscript.

### Isolation of mature rRNA and r-proteins

TP30 and mature 16S rRNA were prepared as described by Nierhaus (15). TP30 was purified using acetone precipitation.

### Expression of 30S and 50S r-proteins in PURE system

Each protein was expressed in a 25 μl PURE system reaction at 37°C for 2 hr with 250 ng plasmid templates. Reactions were later analyzed on 4-12% Bis-Tris Gels (Life Technologies) and stained by Coomassie Blue G-250.

### Preparation of in vitro transcribed 16S rRNA and in vitro synthesized 30S r-proteins

The 16S rRNA gene from *E. coli* MG1655 strain rrnB operon was cloned under the control of a T7 promoter, transcribed in AmpliScribe T7-Flash Transcription Kit (Epicentre) and purified by phenol/chloroform extraction and ethanol precipitation. 30S r-proteins were expressed in a 200 μl PURE system reaction and purified using reverse his-tag purification method (NEB PURE system handbook). Specifically, after expression, each reaction was diluted with 200 μl dilution buffer (50 mM Hepes-KOH, 10 mM Mg acetate, 0.8 M NaCl, 0.1% Tween20) and applied to Amicon Ultracel 0.5 ml-100 K spin concentrator and centrifuged for 50 min at 10000 × g at 4°C. The permeate was transferred to a new tube and mixed with 0.25 volumes Ni-NTA magnetic beads (Qiagen) for 40 min at 4°C. Supernatant was concentrated by Amicon Ultracel 0.5 ml-3 K spin concentrator and exchanged to a storage buffer to a final concentration of 50 mM Tris-HCl pH 7.6, 100 mM KCl, 1 mM DTT and 30% glycerol.

### Integrated 16S rRNA modification, cofactor facilitated ribosome assembly, in vitro transcription and translation

Integrated assay was set up to 15 μl with 6 μl solution A, 1.8 μl factor mix from PURE Δ ribosome kit, 0.8 U/μl Murine RNase Inhibitor (New England Biolabs), 10 ng/μl pIVEX 2.3d Fluc plasmid, 0.3 μM 50S subunit, 0.36 μM or 0.9 μM TP30, 0.3 μM native or *in vitro* transcribed 16S rRNA, 80 μM S-Adenosyl methionine (SAM), various concentrations of 16S rRNA modification enzymes and 30S ribosome assembly cofactors as indicated. When reconstituting fully synthetic 30S subunits, TP30 was replaced by 0.9 μM PURE system synthesized 30S r-proteins and optimized concentrations of 30S ribosome assembly cofactors and 16S rRNA modification enzymes were used. Reactions were incubated at 37°C for 2 hr. After incubation, 7 μl reaction was mixed with 40 μl luciferase assay substrate (Promega) and incubated at room temperature for 10 min. Luminescence was measured by SpectraMaxM5 plate reader (Molecular Devices, Sunnyvale, CA). 5 replicates were conducted for each condition.

### Co-expression of 30S and 50S r-proteins

For 30S subunit, 50 ng of each plasmid encoding r-proteins, 0.8 U/μl Murine RNase Inhibitor (New England Biolabs), 0.3 mM ^13^C labelled Lys, Arg and 0.3 mM other 18 amino acids were added to 100 μl PURE Δ amino acids system reaction and incubated at 37°C for 2 hr. For 50S subunit, 50 ng of each plasmid encoding r-proteins, 0.8 U/μl Murine RNase Inhibitor (New England Biolabs), 0.3 mM ^13^C labelled Lys, Arg and 0.3 mM other 18 amino acids were added to 150 μl PURE Δ amino acids system reaction and incubated at 37°C for 2 hr.

### Mass spectrometry sample preparation

30 μl of co-expression reaction was taken and diluted to 1 mg/ml with Alkylation buffer (8 M Urea, 25 mM Tris-HCl pH 8.0, 10 mM DTT) and incubate at 56°C for 30 min, then cool down to room temperature. Iodoacetamide was added to the protein solution to a final concentration of 30 mM. The tube was wrapped with foil and incubated at room temperature for 30 min. DTT was added to 20 mM and incubated at 37°C for 30 min to quench the reaction. 1/4 volume of 100% TCA was added drop wise to each sample and incubated for 10 min on ice. Each sample was then spun at ~14K rpm for 5 min at 4°C. Supernatant was discarded. Pellet was washed with 200 μl ice cold acetone per tube by vortexing and spun again at ~14K rpm for 5 min at 4°C. The washing step was repeated twice. Pellet was air-dried on bench. 500 μl digestion buffer (8 M Urea, 50 mM Tris-HCl pH 8.0) was added to protein pellet and incubated at 56°C for 60 min to denature the proteins. Each sample was then cooled. 100 mM Tris-HCl (pH 8.0) was added to each sample until urea concentration was less than 1M. 20 μg trypsin was added to protein samples and incubated at 37°C for 6 hr. 1/2 volume of 5% acetonitrile/5% formic acid was added to samples to reach a final pH <4. Water’s tc18 column was pre-wet with 1 ml 100% acetonitrile and 1 ml 90% acetonitrile/5% formic acid and then equilibrated with 1 ml 5% acetonitrile/5% formic acid. Acidified samples were loaded to the column slowly (< 1ml/min), washed with 1 ml 5% acetonitrile/5% formic acid and eluted with 500 μl 50% acetonitrile/ 5% formic acid. Samples were then dried using Speedvac.

### Mass spectrometry data analysis

The generated peptides were analyzed using liquid-chromatography tandem mass spectrometry (LC-MS/MS) essentially as described previously (16). Briefly, the analysis was performed using an Orbitrap Elite mass spectrometer (Thermo Scientific, San Jose, CA) equipped with an Accela 600 binary HPLC pump (Thermo Scientific) and a Famos autosampler (LC Packings). Peptides were fractionated over a 100 μm I.D. in-house-made microcapillary column packed first with approximately 0.5 cm of Magic C_4_ resin (5 μm, 100 Å, Michrom Bioresources) followed by 20 cm of Maccel C_18_AQ resin (3 μm, 200 Å, Nest Group). Fractionation was achieved by applying a gradient from 10 % to 32 % acetonitrile in 0.125 % formic acid over 75 min at a flow rate of approximately 300 nl min^-1^. The mass spectrometer was operated in a data-dependent mode collecting survey MS spectra in the Orbitrap over an m/z range of 300-1500, followed by the collection of MS2 spectra acquired in the dual pressure linear ion trap on the up to 20 most abundant ions detected in the survey MS spectrum. The settings for collecting survey MS spectra were: AGC target, 1x10^6^; maximum ion time, 50 ms; resolution 6x10^3^. The settings for the acquisition of MS2 spectra were: isolation width, 2 m/z; AGC target, 2x10^3^; max. ion time, 100 ms; normalized collision energy, 35; dynamic exclusion, 40 s at 10 ppm.

Peptides were identified from the MS2 data using the Sequest algorithm (17) operated in an inhouse-developed software suite environment that was also applied for filtering the search results and extracting the quantitative data. The data was searched against a protein sequence database comprised of the sequences of *E. coli* ribosomal proteins, of known contaminants such as porcine trypsin, and of 500 nonsense protein sequences derived from all *S. cerevisiae* protein sequences using a fourth order Markov chain model (18). To this forward (target) database component we added a reversed (decoy) component including all listed protein sequences in reversed order (19). The Markov chain model derived nonsense protein sequences were added to generate a protein sequence database size allowing an accurate estimation of the false discovery rate of assigned peptides and proteins. Searches were performed using a 50 ppm precursor ion mass tolerance and we required that both termini of the assigned peptide sequences were consistent with trypsin specificity allowing two missed cleavages. Carbamidomethylation of cysteines (+57.02146) was set as static information and full ^13^C labeling on arginine and lysine (+6.020129) as well as oxidation on methionine (+15.99492) were set as variable modifications. A false discovery rate of less than 1 % for the assignments of both peptides and proteins was achieved using the target-decoy search strategy (19). Assignments of MS2 spectra were filtered using linear discriminant analysis to define one composite score from the following spectral and peptide sequence specific properties: mass accuracy, XCorr, ΔCn, peptide length, and the number of missed cleavages (20). Protein identifications were filtered based on the combined probabilities of being correctly assigned for all peptides assembled into each protein (20). Both peptide and protein assignments were filtered to a false-discovery rate of less than 1 %. Relative peptide quantification was done in an automated manner by producing extracted ion chromatograms (XIC) for the light (^12^C arginine or lysine) and heavy (^13^C arginine or lysine) forms of a peptide ion followed by measuring the area under the chromatographic peaks (21). A peptide ion was considered to be quantified when the sum of the signal-to-noise ratio of both light and heavy form was larger or equal to 10. The median of the log2 (heavy/light) ratios measured for all peptides assembled into a protein was defined as the protein log2 ratio. If only the ion signal of the light peptide form was observed above the noise level the log2 heavy-to-light ratio was defined as being smaller than log2 of the signal-to-noise for the light peptide ion. As orthogonal approach to determine the occurrence of the light and heavy forms of a protein we counted the number of MS2 spectra assigned to each forms of a protein.

## RESULTS

### 30S subunit reconstitution in PURE Δ ribosome system with no supplements

No one, to our knowledge, has integrated ribosome reconstitution and protein synthesis in a well-defined cell free system under physiological conditions. The PURE Δ ribosome system, a ribosome-free version of the PURE system (9) containing defined components required by transcription and translation, can serve as a platform to integrate ribosome reconstitution and measurement of ribosome activity by transcription and translation of a reporter gene in one compartment. As a preliminary test, we reconstituted 30S subunits in the PURE Δ ribosome system with Fluc as a reporter at 37°C in 15 μl reactions. Specifically, 30S subunit reconstitution, its coupling with native 50S subunit, Fluc transcription and translation occurred simultaneously in a 2-hr batch reaction and ribosome activity was measured by quantifying active Fluc synthesized.

We first reconstituted 30S subunits from TP30 and native 16S rRNA or *in vitro* transcribed 16S rRNA. When coupled with native 50S subunits, native 30S subunits exhibited 90% activity of intact native 70S ribosomes, indicating that the recoupling process is not a bottleneck for ribosome reconstitution in our system. Reconstituted ribosomes from native 16S rRNA and TP30 were ~40% as active as native 70S ribosomes, possibly due to lack of assembly cofactors and natural rRNA processing pathways. Reconstituted ribosomes from *in vitro* transcribed 16S rRNA and TP30 showed another ~ 12% drop of activity possibly due to lack of 16S rRNA modifications (Figure 2).

**Figure 2.**
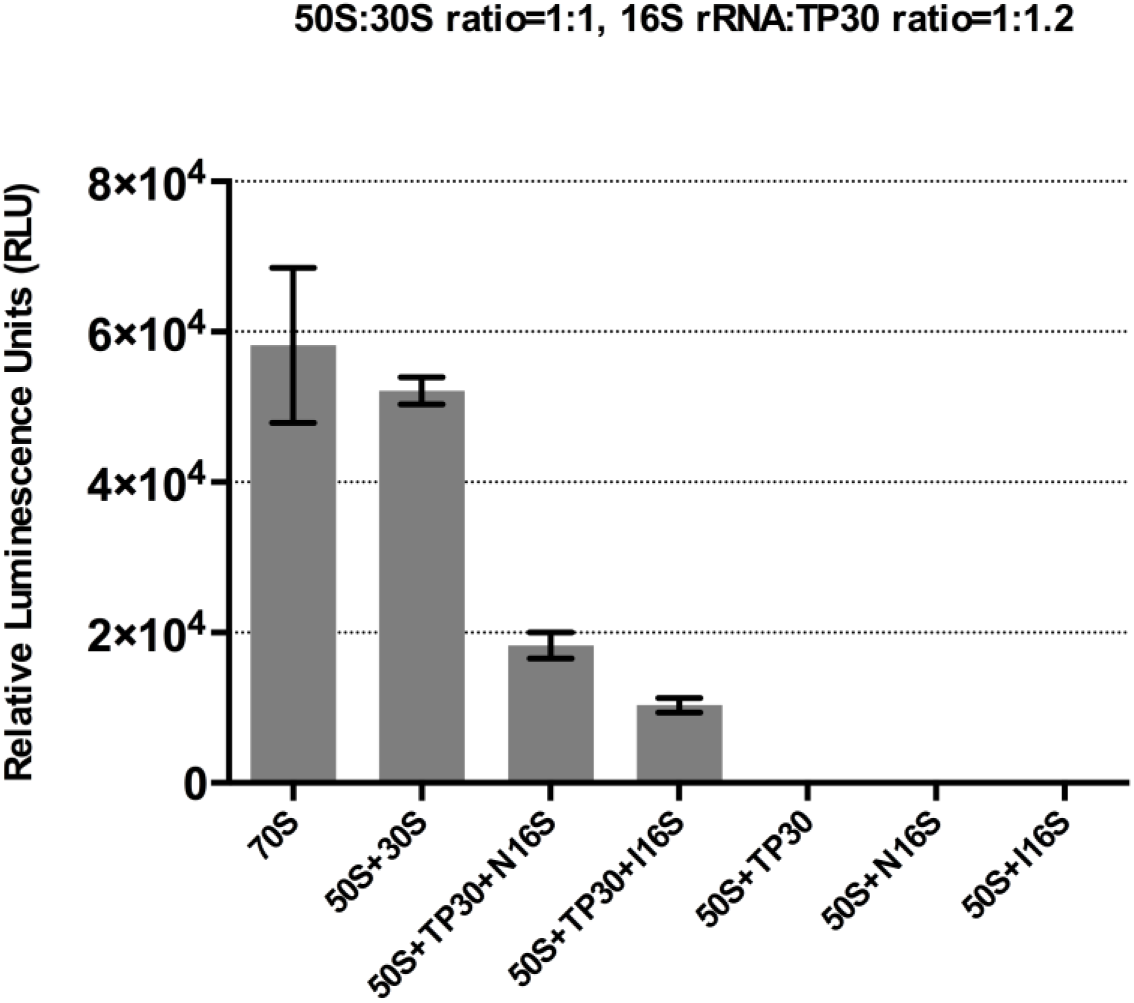
Integrated ribosome assembly, transcription and translation in PURE Δ ribosome system under physiological conditions with TP30 and native or *in vitro* transcribed 16S rRNA. N16S: native 16s rRNA. IVT 16S: *in vitro* transcribed 16S rRNA. For reconstitution reactions, 0.3 μM 70S, 0.3 μM 50S, 0.3 μM 30S, 0.36 μM TP30, 0.3 μM native 16S rRNA, 0.3 μM *in vitro* transcribed 16S rRNA were added as indicated and incubated at 37°C for 2 hr. Intact 70S was taken as a positive control. Luciferase activities were measured in relative luminescence unit. Error bars are ± standard deviations, with n=5.

**Figure 3.**
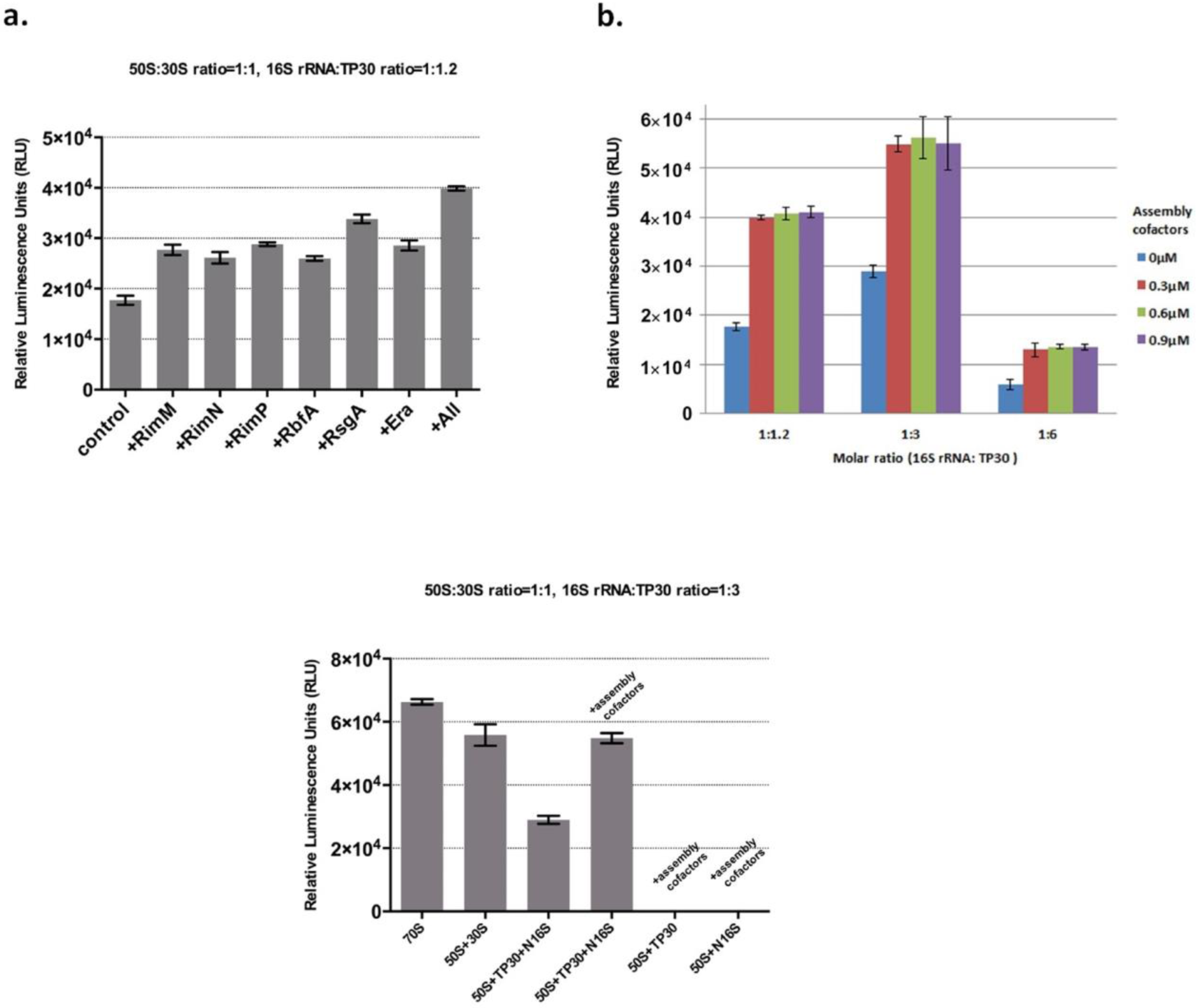
Supplementing 30S ribosome assembly cofactors to 30S subunit reconstitution using our integrated method. **(a)**. Effect of ribosome assembly cofactors on ribosome reconstitution using our integrated method under physiological conditions with native 16S rRNA and TP30 as measured by luciferase production. For each reconstitution reaction, 0.3 μM 50S ribosome, 0.3 μM native 16S rRNA, 0.36 μM TP30 and 0.3 μM ribosome assembly cofactor were added. Reactions were incubated at 37°C for 2 hr. **(b).** Optimization of 16S rRNA: TP30 molar ratio and ribosome assembly cofactor concentration on ribosome reconstitution using our integrated method under physiological conditions with native 16S rRNA and TP30 as measured by luciferase production. **(c).** Ribosome reconstitution from natural components with ribosome assembly cofactors. N16S: native 16S rRNA. For reconstitution reactions, 0.3 μM 70S, 0.3 μM 50S, 0.3 μM 30S, 0.3 μM native 16S rRNA, 0.9 μM TP30 and 0.3 μM of each ribosome assembly cofactor were added as indicated. Reactions were incubated at 37°C for 2 hr. Luciferase activities were measured in relative luminescence unit. Error bars are ± standard deviations, with n=5.

### 30S subunit reconstitution from natural components in PURE Δ ribosome system with 30S ribosome assembly cofactors

We hypothesized that key components of ribosome biogenesis pathway were missed in *in-vitro* ribosome assembly and mimicking ribosome biogenesis *in vivo* would promote ribosome reconstitution. Actually, the rate of ribosomal subunit biogenesis *in vivo* makes it difficult to isolate assembly intermediates and test specific factors that facilitate the process. Using our platform, we were able to assess various ribosome biogenesis factors on ribosome reconstitution *in vitro*. Here we first focused on 30S ribosome assembly cofactors. 30S ribosome assembly cofactor RimM, RimN, RimP, RbfA, RsgA and Era were cloned, purified and analyzed on SDS-PAGE (Supplementary Figure S1). Figure 2a shows RimM, RimN, RimP, RbfA, RsgA and Era, when added at a 1:1 molar ratio to ribosomes, improved reconstituted ribosome activity by ~57%, 49%, 60%, 48%, 94%, 58% respectively, using our integrated method under physiological conditions with native 16S rRNA and TP30. Adding all of them (each at 1:1 molar ratio to ribosome) increased reconstituted ribosome activity by ~2.3 fold.

We subsequently carried out a series of optimization experiments to increase combined ribosome assembly and protein synthesis activity, focusing on 16S rRNA: TP30 molar ratio and the concentration of 30S ribosome assembly cofactors. Results showed that maximum increase in reconstituted ribosome activity was achieved at a 16S rRNA: TP30 molar ratio of 1:3 and a 30S ribosome assembly cofactor mix concentration of 0.3 μM. Further increases of 30S ribosome assembly cofactor mix concentration did not increase reconstituted ribosome activity. 30S subunits assembled with a 16S rRNA: TP30 molar ratio of 1:6 had ~20% the activity of those assembled with a 16S rRNA: TP30 molar ratio of 1:3 (Figure 2b). Reconstituted ribosomes from native 16S rRNA and TP30 with optimized 16S rRNA: TP30 molar ratio had ~40% of native 70S ribosome activity and when combined with optimized 30S ribosome assembly cofactor mix concentration showed an activity of ~90% of native 70S ribosomes (Figure 2c) which is 5-fold better than what Jewett et al achieved using iSAT method in crude *E. coli* S150 extracts which may contain not only those assembly cofactors, but also proteases or nucleases (8) (Table 3).

**Table 3.** Number of peptide bonds produced per ribosome as measured by Fluc synthesized in cell-free systems. (A30S: assembled 30S from native 30S r-proteins and native 16S rRNA. I30S: assembled 30S from native 30S r-proteins and *in vitro* transcribed 16S rRNA. S30S: assembled 30S from *in vitro* synthesized 30S r-proteins and *in vitro* transcribed 16S rRNA.)

| Number of peptide bonds produced per ribosome | iSAT method in *E. coli* S150 extract (8) | PURE system (this study) | PURE system with supplements |
| --- | --- | --- | --- |
| 70S | ~27.5 | ~10 | ~50 (25) |
| 50S+A30S | ~2.2 | ~3.3 | ~10 (this study) |
| 50S+I30S | ~2.2 | ~1.6 | ~6.4 (this study) |
| 50S+S30S | N/A | ~0.64 | ~1.3 (this study) |

### 30S subunit reconstitution from TP30 and in vitro transcribed 16S rRNA in PURE Δ ribosome system with 16S rRNA modification enzymes and 30S ribosome assembly cofactors

We next asked whether 16S rRNA modification could be incorporated into our integrated method. We cloned the eleven 16S rRNA modification enzymes, and purified and analyzed them on SDS-PAGE (Supplementary Figure S2). When supplementing our system with 16S rRNA modification enzymes, we also added 80 μM SAM to the reconstitution reaction as a methyl donor for rRNA methylation. Figure 4a shows ribosome reconstituted from *in vitro* transcribed 16S rRNA with no modifications was only ~40% active as those reconstituted with native 16S rRNA when a 16S rRNA: TP30 molar ratio of 1:1.2 was used. After adding modification enzymes, the reconstituted ribosome activity increased and reached a maximum boost of ~30% at 1.5 μM. Although modification enzymes were added to the reconstitution reaction, it may be that, either individually or as a group, they were unable to modify their corresponding positions on the 16S rRNA at levels comparable to *in vivo* modification. However, the ~30% boost suggests that some modifications were made successfully.

**Figure 4.**
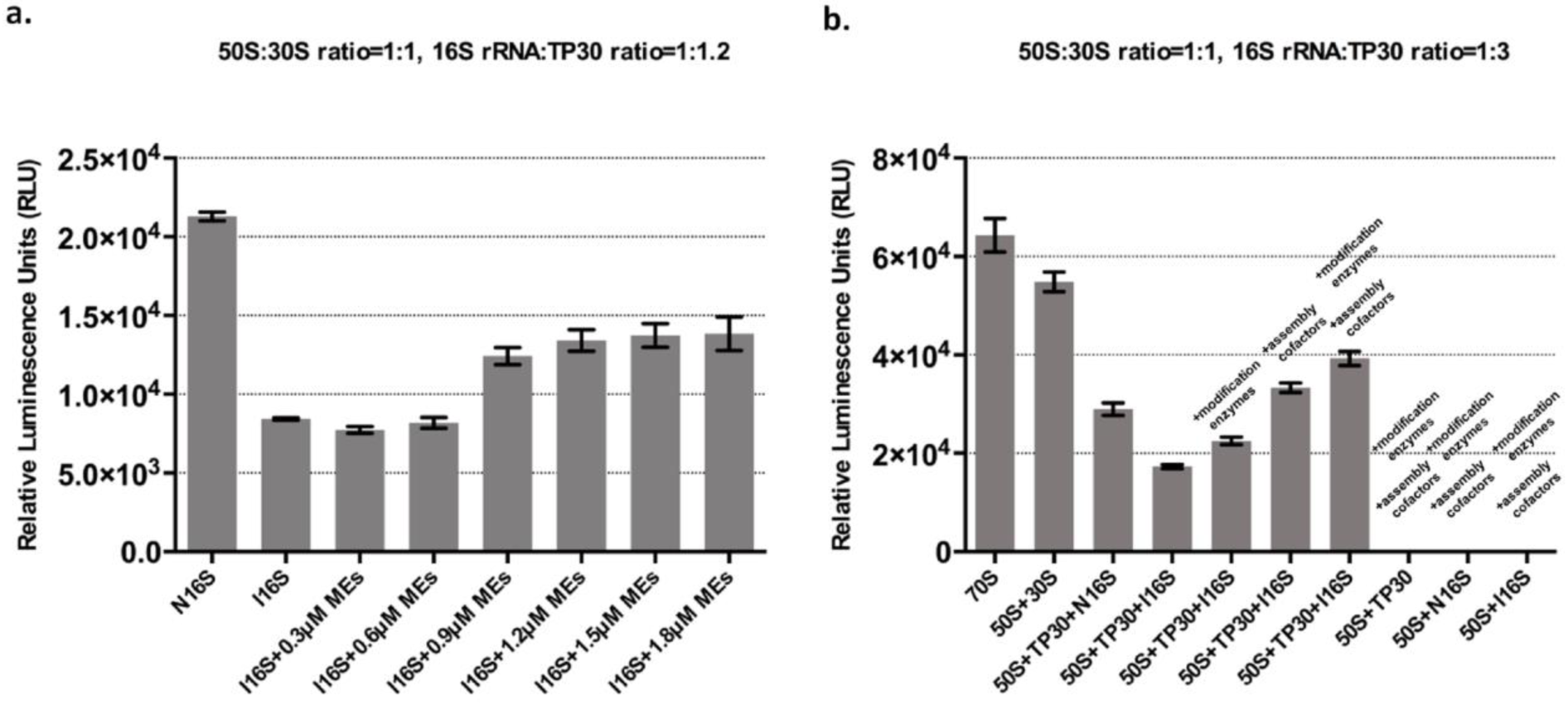
Reconstitute 30S subunits with *in vitro* transcribed 16S rRNA using our integrated method in PURE Δ ribosome system under physiological conditions by supplementing 16S rRNA modification enzymes and 30S ribosome assembly cofactors. (a). 30S ribosome reconstitution with different concentrations of 16S rRNA modification enzymes with *in vitro* transcribed 16S rRNA and TP30 as measured by luciferase production. For each reconstitution reaction, 0.3 μM 50S ribosome, 0.3 μM native or *in vitro* transcribed 16S rRNA, 0.36 μM TP30 and the indicated amount of 16S rRNA modification enzymes were added. Reactions were incubated at 37°C for 2 hr. **(b).** 30S ribosome reconstitution from TP30 and native or *in vitro* transcribed 16S rRNA with optimized concentrations of 16S rRNA modification enzymes and 30S ribosome assembly cofactors as measured by luciferase production. N16S: native 16s rRNA. I16S: *in vitro* transcribed 16s rRNA. For reconstitution reactions, 0.3 μM 70S, 0.3 μM 50S, 0.3 μM 30S ribosomes, 0.3 μM native or *in vitro* transcribed 16S rRNA, 0.9 μM TP30, 0.3 μM of each ribosome assembly cofactor and 1.5 μM of each 16S rRNA modification enzymes were added as indicated. Reactions were incubated at 37°C for 2 hr. Luciferase activities were measured in relative luminescence unit. Error bars are ± standard deviations, with n=5.

To assess if combining 30S ribosome assembly cofactors and 16S rRNA modification enzymes was beneficial, we assembled 30S subunits from *in vitro* transcribed 16S rRNA and TP30 in the PURE Δ ribosome system supplemented with 30S ribosome assembly cofactors and 16S rRNA modification enzymes at their optimized concentrations. Here the optimized 16S rRNA: TP30 molar ratio 1:3 was used. Ribosomes reconstituted from *in vitro* transcribed 16S rRNA and TP30 with 16S rRNA modification enzymes had ~72% activity of ribosomes reconstituted from native 16S rRNA and ~33% activity of native 70S ribosomes. Ribosomes reconstituted from *in vitro* transcribed 16S rRNA and TP30 with 30S ribosome assembly cofactors had ~55% activity of native 70S ribosomes. When 30S ribosome assembly cofactors and 16S rRNA modification enzymes were combined, we were able to assemble functional ribosomes with ~70% activity of native 70S ribosomes from *in vitro* transcribed 16S rRNA and TP30 (Figure 4b).

### 30S subunit reconstitution from in vitro synthesized 30S r-proteins and in vitro transcribed 16S rRNA in PURE Δ ribosome system with 16S rRNA modification enzymes and 30S ribosome assembly cofactors

The successful reconstitution of active 30S subunits with *in vitro* transcribed 16S rRNA has been previously reported (3), but to our knowledge this has never been done with *in vitro* synthesized 30S r-proteins instead of TP30. Thus, in an effort to make fully synthetic 30S subunits, we first cloned all genes encoding 30S r-proteins into pET-24b vectors and expressed them in PURE system (Supplementary Figure S3). We then purified each protein using the reverse his-tag purification method described in PURE system handbook. Each purified protein was analyzed by SDS-PAGE (Supplementary Figure S4) and concentrations were determined by Bradford assay. Finally, we replaced TP30 with PURE system synthesized 30S r-proteins in our integrated reconstitution reaction. Here optimized 16S rRNA: r-protein molar ratio and optimized concentrations of 30S assembly cofactors and 16S rRNA modification enzymes were used. Reconstituted 30S subunits from PURE system synthesized 30S r-proteins and native 16S rRNA showed ~12% activity of native ribosomes, which increased to ~21% when supplemented with 30S ribosome assembly cofactors. Fully synthetic 30S subunits assembled from PURE system synthesized 30S r-proteins and *in vitro* transcribed 16S rRNA had ~8% of native ribosome activity, which increased to ~10% when supplemented with 30S ribosome assembly cofactors and ~17% when supplemented with 30S ribosome assembly cofactors and 16S rRNA modification enzymes (Figure 5).

**Figure 5.**
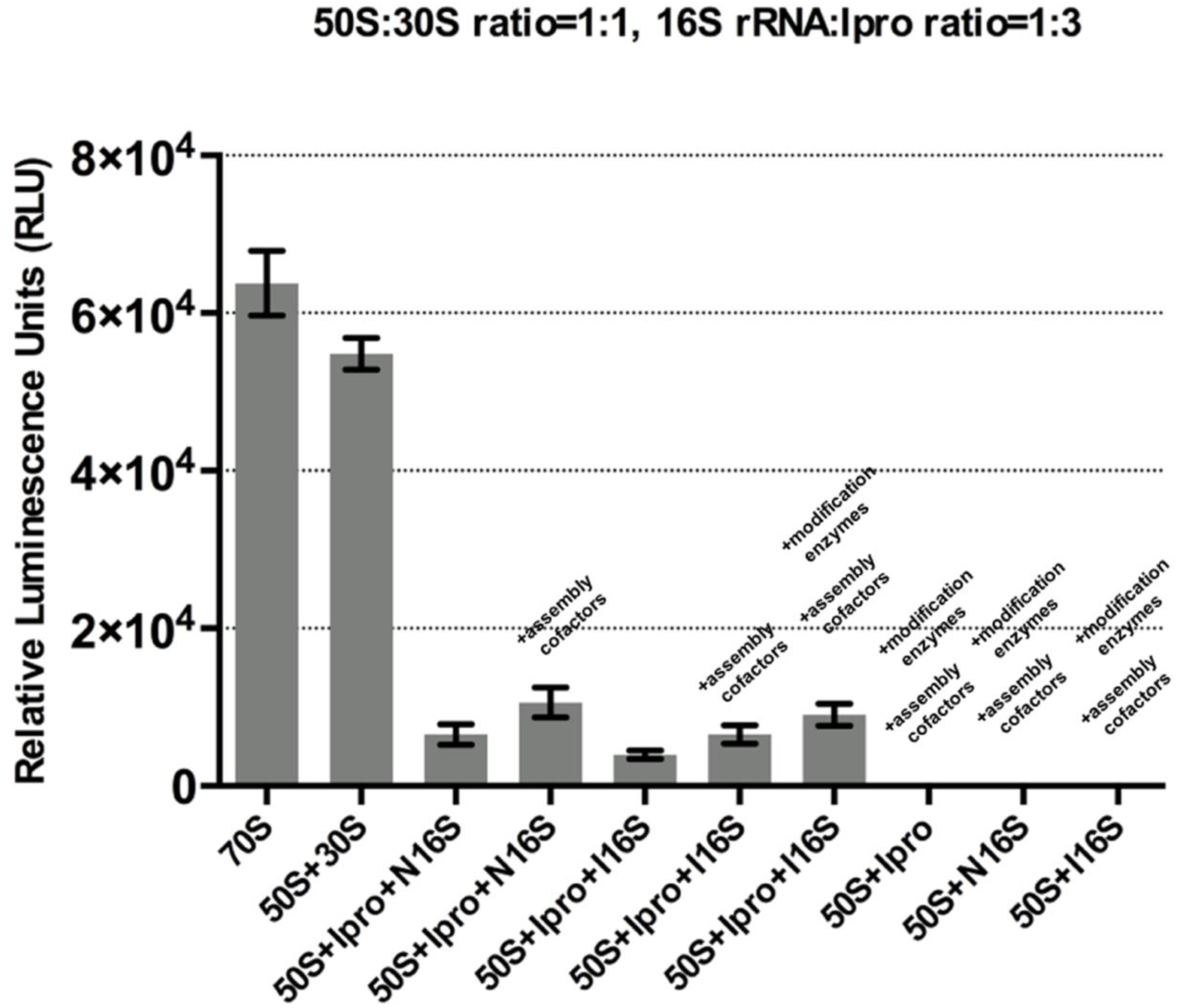
Reconstitute 30S subunits with *in vitro* synthesized 30S r-proteins using our integrated method in PURE Δ ribosome system under physiological conditions by supplementing 30S ribosome assembly cofactors and 16S rRNA modification enzymes. N16S: native 16s rRNA. IVT 16S: *in vitro* transcribed 16S rRNA. Ipro: *in vitro* synthesized 30s r-proteins. For reconstitution reactions, 0.3 μM 70S, 0.3 μM 50S, 0.3 μM 30S ribosomes, 0.3 μM native or *in vitro* transcribed 16S rRNA, 0.9 μM *in vitro* synthesized 30S r-proteins, 0.3 μM of each ribosome assembly cofactor and 1.5 μM of each 16S rRNA modification enzymes were added as indicated. Reactions were incubated at 37°C for 2 hr. Luciferase activities were measured in relative luminescence unit. Error bars are ± standard deviations, with n=5.

## DISCUSSION

In this study, we followed a strategy of mimicking native ribosome biogenesis pathways in a well-defined cell free system in which ribosomes could be assembled under physiological conditions, and successfully integrated rRNA modification, ribosome assembly, and *in vitro* transcription and translation by supplementing it with corresponding assembly cofactors and modification enzymes. In this system, we could study and optimize 30S subunit construction using many combinations of natural and synthetic parts.

Table 3 compares productivity of native, reconstituted semi-synthetic and fully synthetic ribosomes in different cell free systems as measured by number of peptide bonds produced per ribosome. Compared to iSAT method in crude *E. coli* S150 extracts (8), our method (with supplements) achieved ~5 fold higher productivity for assembled 30S subunits from native 16S rRNA and TP30. We suspect that it is mainly because in our well-defined system, we could easily control assembly cofactor concentrations to achieve highest ribosome reconstitution efficiency while avoiding protease and nuclease contamination. In fact, our system demonstrates higher engineering flexibility than crude extract based systems by allowing screening or testing of ribosome biogenesis factors, not only assembly cofactors, on their functions on ribosome assembly.

Our method (with supplements) achieved ~3 fold higher productivity than iSAT method (8) for assembled 30S subunits from *in vitro* transcribed 16S rRNA and TP30 (Table 3). We showed that introducing 16S rRNA modification enzymes during ribosome assembly increased reconstituted ribosome activity. In fact, these enzymes modify 30S at different assembly stages (Table 2). In our platform, ribosome assembly can proceed through all stages in a single compartment, allowing modification enzymes to find their ideal substrates. Monitoring the progress of assembly in our system should therefore offer new opportunities to study rRNA modification dynamics *in vitro*. We note that even with the addition of ribosome assembly cofactors and 16S rRNA modification enzymes, the *in vitro* transcribed 16S rRNA may still not be properly folded because the rRNA is not produced in its natural pathways (22). This suggests a possible follow-on in which we attempt to enable rRNA processing pathways in our system by supplementing it with endonucleases and helicases.

This paper also documents the first construction of functional active 30S subunits with *in vitro* transcribed 16S rRNA and *in vitro* synthesized r-proteins. With synthetic proteins and rRNA, the reconstitution efficiency decreased to ~17% of native ribosome activity even with the presence of 30S ribosome assembly cofactors and 16S rRNA modification enzymes. Besides the missing natural rRNA processing pathway and the incompleteness of rRNA modification just discussed, additional hypotheses for explaining this low efficiency come to mind: First, r-proteins expressed in PURE system may not be as well folded as *in vivo* which could lead to a significant decrease in the reconstituted ribosome activity. Second, in living cells, some r-proteins are modified after translation, such as S5, S6, S11, S12 and S18 (23), potentially affecting ribosome activity or translation fidelity. The functions of these modifications are not well understood yet, but can be characterized using our platform in the near future.

Given the success of our strategy of mimicking ribosome biogenesis pathway *in vivo* with defined biochemical factors for 30S subunit construction, we are aiming to similarly construct synthetic 50S subunits as the next step. This integrated method can potentially be applied to construct fully synthetic 50S subunits and study the effects of 50S ribosome assembly cofactors and 23S rRNA modification enzymes on ribosome biogenesis. As initial steps in this direction, we have demonstrated that 50S r-proteins can be synthesized in the PURE system (Supplementary Figure S5).

Further, we also co-expressed 30S and 50S r-proteins in the PURE system containing ^13^C-labeled arginine and lysine allowing us to quantify their expression by applying a quantitative mass spectrometry-based proteomics approach similar to SILAC (24) (Supplementary Table S8 and S9). We found that 19 of the 21 30S r-proteins as well as 19 out the 33 50S r-proteins were identified as carrying ^13^C-labeled lysine or arginine, showing that these proteins were successfully co-expressed in our system. The ratio of newly synthesized protein to protein of PURE system ribosomes was found to as high as 1:1 (RpsK) for 30S proteins and 1:4 (RplP) for 50s proteins showing high efficiency of the expression. Further optimization of PURE system will lead to higher yields of r-proteins (10,25-28). We plan to explore balancing of protein expression levels by using codon optimization and by adjusting the amount of DNA templates added.

Making functional fully synthetic ribosomes under physiological conditions is considered one of the main barriers to construct a self-replicating entity from simple biochemical components (11, 12). Here we have taken significant steps towards fully synthetic ribosomes in a system consisting solely of purified components by integrating rRNA modification, cofactor facilitated ribosome assembly and protein synthesis together. We are the first group to document the construction of functional fully synthetic 30S subunits. We extended the minimal genome proposed in (11) of 151 gene products to at least 180 for fast and accurate macromolecular synthesis. Further improvements of the system may allow us to construct the proposed DNA, RNA and protein based system capable of central-dogma replication.

## SUPPLEMENTARY DATA

Supplementary Data are available at NAR Online.

## ACKNOWLEDGEMENTS

We thank Dr. Poyi Huang, Dr. Liangcai Gu, Dr. Anthony Forster, and Dr. Ben Stranges for advice and discussions, A Joachimiak for sharing pET 15b-yhhF, M Deutscher for sharing pET-32a yggJ, K Rudd for sharing pET-28a yhiQ.

## AUTHOR CONTRIBUTIONS

JL and GMC designed the experiments; JL and BW did the experiments. All authors helped draft and edit the final manuscript.

